# Hierarchical Computational Anatomy: Unifying the Molecular to Tissue Continuum Via Measure Representations of the Brain

**DOI:** 10.1101/2021.04.19.440540

**Authors:** Michael Miller, Daniel Tward, Alain Trouvé

## Abstract

**Objective:** The objective of this research is to unify the molecular representations of spatial transcriptomics and cellular scale histology with the tissue scales of Computational Anatomy for brain mapping.

**Impact statement:** We present a unified representation theory for brain mapping based on geometric measures of the micro-scale phenotypes of molecular disease simultaneously with the connectomic scales of complex interacting brain circuits.

**Introduction:** Mapping across coordinate systems in computational anatomy allows us to understand structural and functional properties of the brain at the millimeter scale. New measurement technologies in digital pathology and spatial transcriptomics allow us to measure the brain molecule by molecule and cell by cell based on protein and transcriptomic identity. We currently have no mathematical representations for integrating consistently the tissue limits with the molecular particle descriptions. The formalism derived here demonstrates the methodology for transitioning consistently from the molecular scale of quantized particles – as first introduced by Dirac as the class of generalized functions – to the continuum and fluid mechanics scales appropriate for tissue.

**Methods:** We introduce two methods based on notions of generalized function geometric measures and statistical mechanics. We use generalized functions expanded to include functional geometric descriptions - electrophysiology, transcriptomic, molecular histology – to represent the molecular biology scale integrated with a Boltzman like procedure to pass from the sparse particles to empirical probability laws on the functional state of the tissue.

**Results:** We demonstrate a unified mapping methodology for transferring molecular information in the transcriptome and histological scales to the human atlas scales for understanding Alzheimer’s disease.

**Conclusions:** We demonstrate a unified brain mapping theory for molecular and tissue scales based on geometric measure representations.

## 1 Introduction

One of the striking aspects of the study of the brain in modern neurobiology is the fact that the distributions of discrete structures that make up physical tissue, from neural cells to synapses to genes and molecules, exists across nearly ten orders of magnitude in spatial scale. This paper focusses on the challenge of building multi-scale representations that simultaneously connect the quantum nano-scales of modern molecular biology and digital pathology for characterizing neural circuits architecture in the functioning brain and disease with the classical continuum representations at the anatomical gross and meso scales.

We have been highly motivated by the Cell Census Network project (BICCN [1]) which brings the nano and micron scales of single cell measures of RNA via spatial transcriptomics [2–4] coupled to the tissue scales of mouse atlases. The recent review on bridging scales from cells to physiology [5] motivates the mathematical framework presented herein. The recent emergence of spatial transcriptomics as Nature method of the year highlights the importance and ascendence of such approaches for understanding the dense metric structure of the brain which represent the coarse physiological atlas scales built up from dense imaging measurements at the cellular scales. Specifically, in our own work digital pathology for the study of Alzheimer’s disease called the BIOCARD study [6], we are examining pathological Tau at both the micro histological and macro atlas scales of Tau particle detections, from 10-100 *μ*m [7, 8] and to human magnetic resonance millimeter scales for examining entire circuits in the medial temporal lobe. In the mouse cell counting project we are examining single-cell spatial transcriptomics using modern RNA sequencing in dense tissue at the micron scale and its representations in the Allen atlas coordinates [9].

Most noteworthy for any representation is that at the finest micro scales nothing is smooth; the distributions of cells and molecules are more well described as random quantum counting processes in space [10]. In contrast, information associated to atlasing methods at the gross anatomical tissue and organ scales of Computational Anatomy extend smoothly [11–16]. Cross-sectionally and even cross-species, gross anatomical labelling is largely repeatable, implying information transfers and changes from one coordinate system to another smoothly. This is built into the representation theory of diffeomorphisms and soft matter tissue models for which advection and transport hold [17–21], principles upon which continuum mechanics and its analogues are based. Also of note is the fact that the brain organizes information on geometric objects, submanifolds of the brains such as the foliation of the cortex and associated coordinates of the cortical columns. Our representations must both represent the quantum to ensemble scales as well as encode the geoemetric organization of the brain.

The focus of this paper is to build a coherent representation theory across scales. For this we view the micron to millimeter scales via the same representation theory called mathematical *geometric measures*, building the finest micron scales from discrete units termed particle measures which represent molecules, synapses and cells. The measure representation from fine to coarse scale aggregates forming tissue. This measure representation allows us to understand subsets of tissues that contain discretely positioned and placed functional objects at the finest quantized scales and simultaneously pass smoothly with aggregation to the classical continuum scales at which stable functional and anatomical representations exist. Since the study of the function of the brain on its geometric submanifolds -the gyri, sulci, subnuclei and laminae of cortex- are so important, we extend our general framework to exploit varifold measures [22] arising in the modern discipline of geometric measure theory. Geometric measures are a class of generalized functions which have the basic measure property of additivity on disjoint unions of the experimental probe space and encode the complex physiological functions with the geometric properties of the submanifolds to which they are associated. To be able to compare the brains we use diffeomorphisms as the comparator tool, with their action representing 3D varifold action which we formulate as “copy and paste” so that basic particle quantities that are conserved biologically are combined with greater multiplicity and not geometrically distorted as would be the case for measure transport.

The functional features are represented via generalized Dirac delta functions at the finest micro structure scales. The functional feature is abstracted into a function space rich enough to accomodate the molecular machinery as represented by RNA or Tau particles, as well as electrophysiology associated to spiking neurons, or at the tissue scales of medical imaging dense contrasts of magnetic resonance images (MRIs). We pass to the classical function continuum via introduction of a scale-space that extends the descriptions of cortical micro-circuits to the meso and anatomical scales. This passage from the quantized features to the stochastic laws is in fact akin to the Boltzman program transferring the view from the Newtonian particles to the stable distributions describing them. For this we introduce a scale-space of kernel density transformations which allows us to retrieve the empirical averages represented by the determinism of the stochastic law consistent with our views of the macro tissue scales.

The representation provides a recipe for scale traversal in terms of a cascade of linear space scaling composed with non-linear functional feature mapping. Following the cascade implies every scale is a measure so that a universal family of measure norms can be introduced which simultaneously measure the disparety between brains in the orbit independent of the probing technology, RNA identities, Tau or amyloid histology, spike trains, or dense MR imagery.

Our brain measure model implies the existence of a sequence. This scale-space of pairs, the measure representation of the brain and the associated probing measurement technologies we call Brainspace. To formulate a consistent measurement and comparison technology on Brainspace we construct a natural metric upon it allowing us to study its geometry and connectedness. The metric between brains is constructed via a Hamiltonian which defines the geodesic connections throughout scale space, providing for the first time a hierarchical representation that unifies micro to millimeter representation in the brain and makes Brainspace into a metric space. Examples of representation and comparision are given for Alzheimer’s histology integrated to magnetic resonance imaging scales, and spatial transcriptomics.

## 2 Results

### 2.1 Measure Model of Brain Structures

To build a coherent theory we view the micron to anatomical scales via the same representation theory building upon discrete units termed particles or atoms. As they aggregate they form tissues. This is depicted in Figure 1 in which the top left panel shows mouse imaging of CUX1 labelling of the inner layers of mouse cortex (white) and CTP2 imaging of the outer layers (green) at 2.5 micron in plane resolution. Notice the discrete nature of the cells clearly resolved which form the layers of tissue which are the global macro scale features of layer 2,3,4 which stain more prolificaly in white and the outer layers 5,6 which stain more prolifically in green.

**Figure 1:**
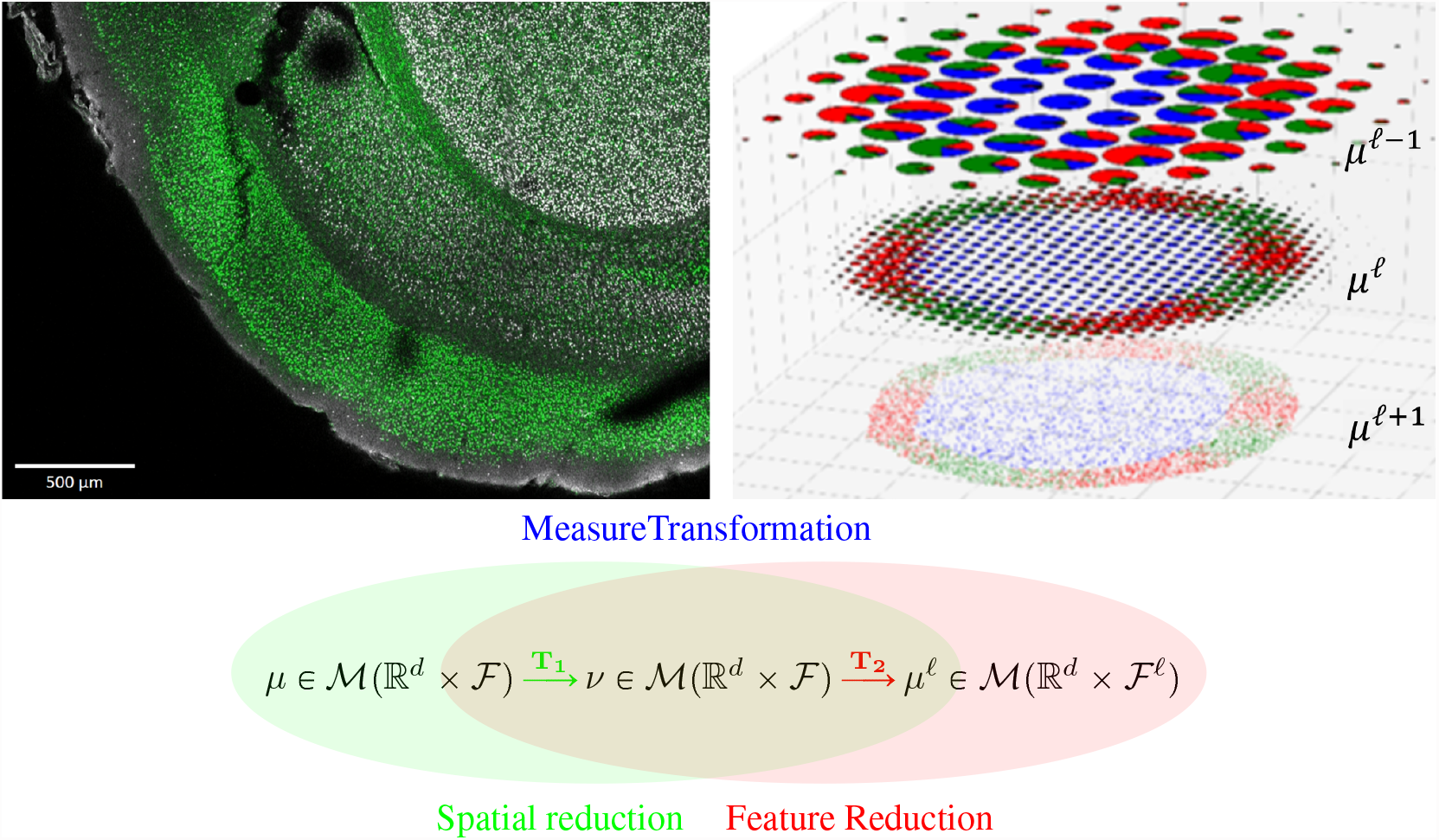
Top Left: Tissue from a NexCre+/-;Brn2fl/+ adult mouse mimicing a wild-type mouse with CUX1 labelling of layer ii/iii,iv and CTIP2 in layers v,vi in green. Shows sections at 2.5^2^ × 50 *μ*m^3^ 6 tile images, 1433 × 1973 pixels; taken from Uli Mueller. Top Right: Showing the abstraction of a coarse-to-fine hierarchy *μ*^𝓁−1^, *μ*^𝓁^, *μ*^𝓁+1^ with fine molecular scales shown at the bottom with colors depicting ℱ function ascending scales. Bottom: Space and function transformation shown as a composition 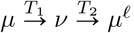.

Our representation exists simultaneously at both the micro and tissue millimeter scales. A key aspect of anatomy is that at a micro or nano scale, information is encoded as a massive collection of pairs (*x*_*i*_, *f*_*i*_) where *x*_*i*_ ∈ ℝ^*d*^ (*d* = 2, 3) describes the position of a “particle” and *f*_*i*_ is a functional state in a given set ℱ attached to it. In our applications ℱ are proteins representing RNA signatures or Tau tangles, and for single cell Neurophysiology represents the dynamics of neural spiking. At the micro scale basically everything is deterministic, with every particle attached to its own functional state among possible functional state in ℱ. But zooming out, the tissue level, say mm scale, appears through the statistical distribution of its constituents with two key quantities, *the local density* of particles *ρ* and the *conditional probability distribution* of the functional features *μ*_*x*_(*df*) at any location *x*. At position *x*, we no longer have a deterministic functional state but a probability distribution *μ*_*x*_ on functional states.

The integration of both descriptions into a common mathematical framework can be done quite naturally in the setting of mathematical measures which are mathematical constructs that are able to represent both the discrete and continuous worlds as well as natural level of approximation between both. We associate the elementary ‘Dirac’ *δ*_*x*_ ⊗ *δ*_*f*_ which applies to infinitesimal volumes in space and function so that *δ*_*x*_ ⊗ *δ*_*f*_ (*dx, df*) = *δ*_*x*_(*dx*)*δ*_*f*_ (*df*) is equal to 1 if *x* ∈ *dx* and *f* ∈ *df* and 0, otherwise. Indeed the set ℳ (ℝ^*d*^ × ℱ) of finite positive measures on ℝ^*d*^ × ℱ contains discrete measures written as

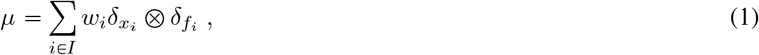

where *w*_*i*_ is a positive weight that can encode the collection (*x*_*i*_, *f*_*i*_) at micro scale.

As in Boltzmann modelling we describe the features statistically at a fixed spatial scale transferring our attention to their stochastic laws modelled as conditional probabilities in ℳ_*P*_ (ℱ) with integral 1. For this we factor the measures into the marginal density measure on space *ρ*^*μ*^ on ℝ^*d*^ with *ρ*^*μ*^(*dx*) = ∫_ℱ_ *μ*(*dx, df*), and the field of probability distributions on ℱ conditioned on *x*. We use the convention *dx* and *df* as events in space and function, respectively, implying the classical factorization

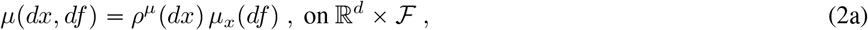

with field of conditional probabilities:

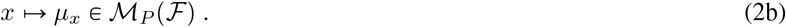

Continuous tissues we abstract as brain measures *μ* with marginal *ρ*^*μ*^ having a tissue density *ρ*(*dx*) = *ρ*_*c*_(*x*)*dx* with respect to the Lebesgue measure on ℝ^*d*^. A fundamental link between the molecular and continuum tissue can be addressed through the law of large numbers since if (*x*_*i*_, *f*_*i*_)_*i* ⩾0_ is an independent and identically distributed sample drawn from law *μ*/*M* of ℝ^*d*^ × ℱ where 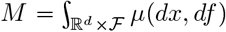 is the total mass of such *μ*, then we have almost surely the weak convergence

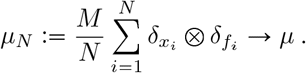

Passing from the tissue scales to the molecular-cellular scales behooves us to introduce a scale-space so that empirical averages which govern it are repeatable. Figure 1 (right) depicts our multi-scale model of a brain as a sequence of measures:

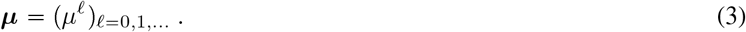

Our idealization of Brainspace as a sequence of measures as depicted in Figure 1 descends from the the coarse tissue scale (top) to the finest particle representation (bottom), with color representing function *f* ∈ ℱ, and radius space-scale. Throughout the range of scales is denoted shorthand 𝓁 < 𝓁_*max*_ to mean 0 ⩽ 𝓁 < 𝓁_*max*_ with lowest scale 𝓁 = 0 and upper 𝓁_*max*_ not attained.

### 2.2 Nonlinear Transformation Model for Crossing Scales

The brain being a multi-scale collection of measures requires us to be able to transform from one scale to another. We do this by associating a scale-space to each particle feature by pairing to each measure a kernel function transforming it from a generalized function to a classical function *δ*_*z*_ : *h* ↦ *h*(*z*). The kernels carry resolution scales *σ* or reciprocally bandwidths, analogous to Planck’s scale.

We introduce the abstract representation of our system as a collection of descriptive elements *z* ∈ *Ƶ* made from spatial and functional features. We transform our mathematical measure *μ*(·) on *Ƶ* generating new measures *μ*′(·) on *Ƶ*′ by defining correspondences via kernels *z* ↦ *k*(*z, dz*′), with the kernel acting on the measures transforming as

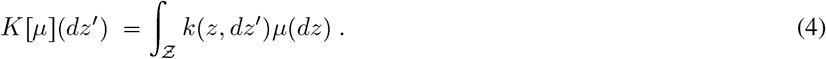

This implies the particles transform as 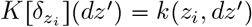.

Figure 1 (bottom row) shows the cascade of operations operations *μ* ↦ *ν* on ℱ transforming linearly, the second *ν* ↦ *μ*^𝓁^ on ℱ^𝓁^ nonlinearly; transforming 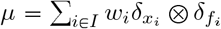 to scale 𝓁 gives

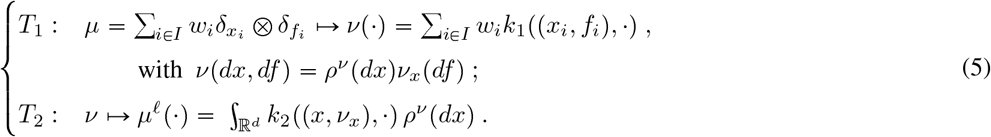

That *T*_1_ : *μ* ↦ *ν* is linear in *μ* follows from 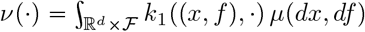.

We use space resampling defined by *π*(*x, y*) the fraction particle *x* shares with *y*, giving density 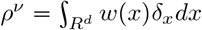 and

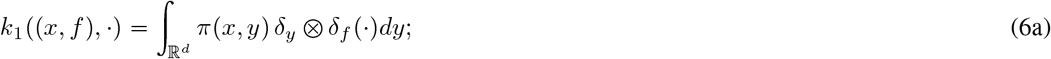

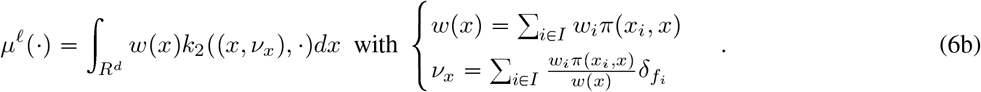

Feature reduction uses singular kernels with projective feature maps from machine learning, *α* ↦ *ϕ*(*α*) ∈ ℱ^𝓁^, *∫*_ℱ_ *α*(*df*) = 1:

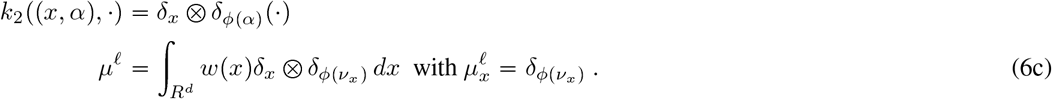

We compute via lattices 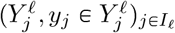 transforming to scale 𝓁 (see Methods 4.2) generating measures on the lattice:

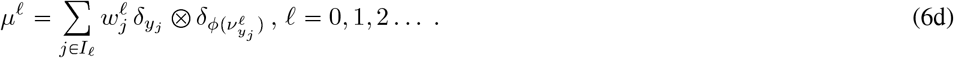

### 2.3 Dynamical Systems Model via Varifold Action of Multi-scale Diffeomorphisms

We want to measure and cluster brains by building a metric space structure. We do this by following the original program of D’Arcy Thompson building bijective crrespondence. In this setting this must be done at every scale with each scale having different numbers of particles and resolutions. We build correspondence between sample brains via dense connections of the discrete particles to the continuum at all scales using the diffeomorphism group and diffeomorphic transport. For this define the group of k-times continuously differentiable diffeomorphisms *φ* ∈ *G*_*k*_ with group operation function composition *φ*^*a*^ ° *φ*^*b*^. For any brain 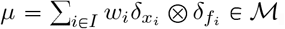, the diffeomorphisms act

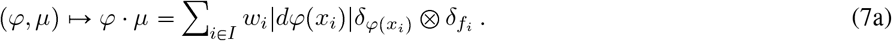

Space scales are represented as the group product, 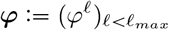, acting component-wise with action

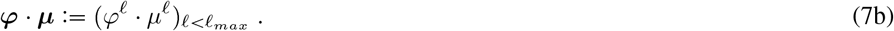

The |*dφ*(*x*)| term in the action we call the “copy and paste” varifold action. It enables the crucial property that when a tissue is extended to a larger area, the total number of its basic constituents increase accordingly with total integral not conserved, in contrast to classic measure or probability transport.

Dynamics occurs by generating the diffeomorphism as flows 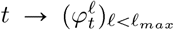, with dynamics controlled by vector fields 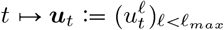 via the ordinary differential equation (ODE) at each scale satisfying

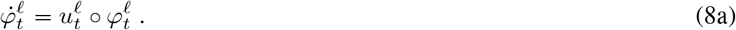

The controls are coupled by successive refinement *v*^𝓁^, 𝓁 *<* 𝓁_*max*_, with *u*^−1^ = 0:

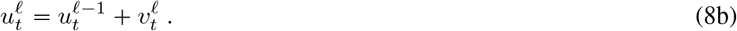

To control smoothness of the maps we force the vector fields to be elements of a reproducing kernel Hilbert spaces (RKHS’s) *V*_𝓁_, norms 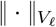, with multi-scale 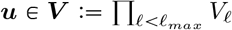. Each RKHS is taken to have a diagonal kernel *K*_𝓁_(·, *·*) = *g*_𝓁_(·, *·*)id_*d*_, with *g*_𝓁_ the Green’s functions with id_*d*_ the *d* × *d* identity (see [24] for non-diagonal kernels). Geodesic mapping flows under a control process along paths of minimum energy respecting the boundary conditions.

Figure 2 shows the multi-scale control hierarchy.

**Figure 2:**
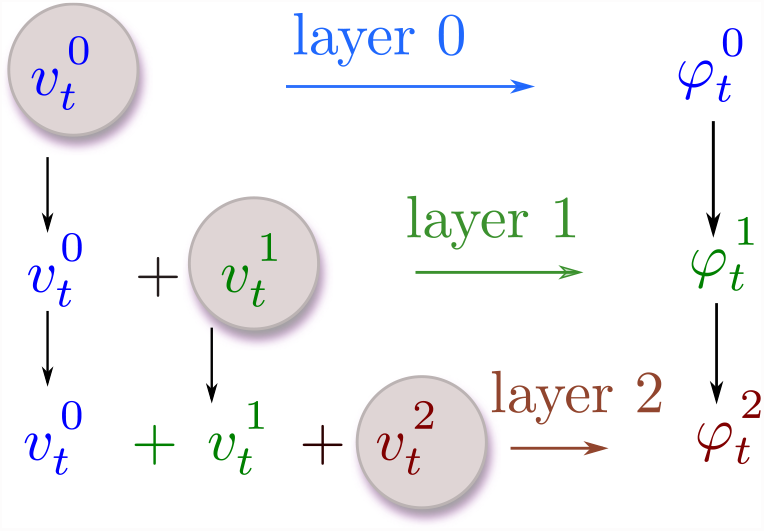
Hierarchical system, controls *u*^𝓁^ = *u*^𝓁−1^ + *v*^𝓁^, *u*^0^ = *v*^0^ and flows *φ*^𝓁^, 𝓁 = 0, 1,….

The multi-scale dynamical control are written 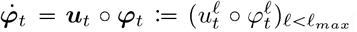. The dynamical system is an observer and dynamics equation:

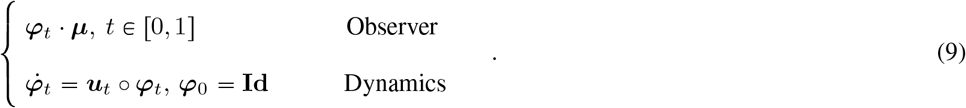

Dynamics translates into a navigation in the orbit of brains and provides a metric distance between brains. Paths of minimum energy connecting the identity ***φ***_0_ = **Id** to any fixed boundary condition (BC) ***φ***_1_ where ***φ***_1_ is accessible defines the distance extending LDDMM [23] to a hierarchy of diffeomorphisms, and is a geodesic for an associated Riemannian metric [24].

The metric from ***μ***_0_ to ***μ***_1_ in the orbit accesible from ***μ***_0_ via diffeomorphisms is the shortest length geodesic paths with BCs ***φ***_0_·***μ***_0_ = ***μ***_0_ and ***φ***_1_·***μ***_0_ = ***μ***_1_. This extension to multi-scale LDDMM Eqn. (23) is given in the Methods 4.4 where we discuss the smoothness required for the geodesics to define a metric and specify the optimal control problem in the state Eqn. (25).

### 2.4 Geodesic Brain Mapping via the Varifold Measure Norm

The BC for matching two brains is defined using measure norms with equality meaning brains are equal, with small normed difference meaning brains are similar. Every brain has a variable number of particles, without correspondence between particles. Measure norms accomodate these variabilities. Geodesic mapping solves for the control minimizing the energy with the boundary endpoint condition modelled via measure norms. The endpoint is a varifold norm constructed from the basic inner product on pairs of points involving a smooth kernel

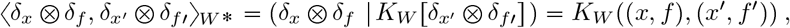

with the kernel defined as separable Gaussians in space and function (see Methods 4.3). The measure norm-square becomes

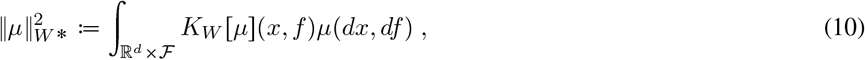

the hierarchical norms across the scales becomes 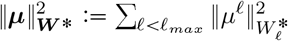.

The optimal control (***u***_*t*_)_0 ⩽ *t* ⩽ 1_ is square-integrable under the *V* -norms, satisfying for *α* > 0:

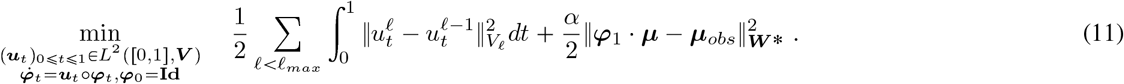

Hamiltonian control parameterizes the measures 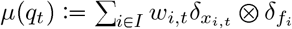 via the flows of the state processes

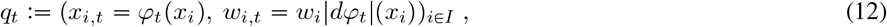

giving endpoint 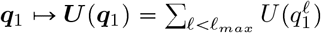 modelled as continuously differentiable in the states:

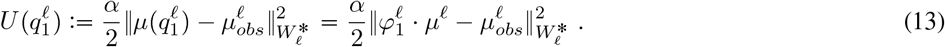

Hamiltonian control reparameterizes (11) in “co-states” 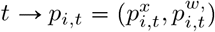, define 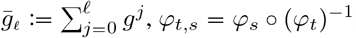:

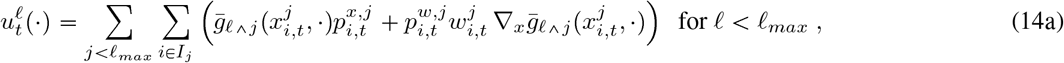

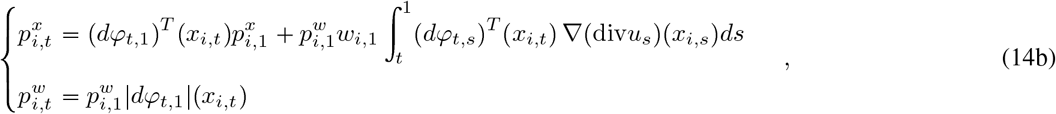

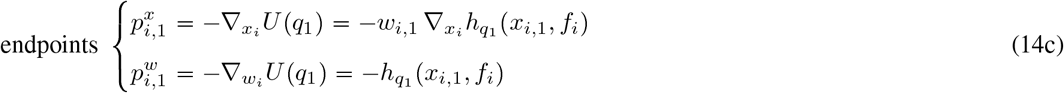

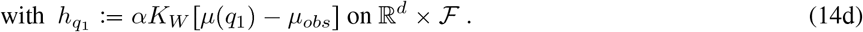

In the above we have eased notation removing indexing of the optimal controls by scales 𝓁 when implied. Methods 4.5 establishes the smoothness for the Hamiltonian equations. Methods 4.6 establishes the smoothness for the norm gradients.

#### 2.4.1 Gradients of the norm endpoints unifying the molecular and tissue models

Calculating the variations on dense voxel images unifies the tissue scales with the sparse molecular scales. Imaging at the tissue continuum scales has the measures as dense limits 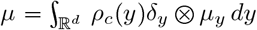 ; defining the state *q*_*t*_ := (*φ*_*t*_, *w*_*t*_ = *w*_0_|*dφ*_*t*_|) gives

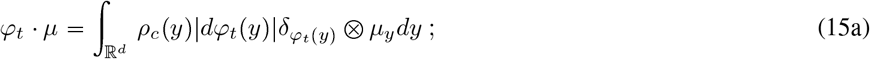

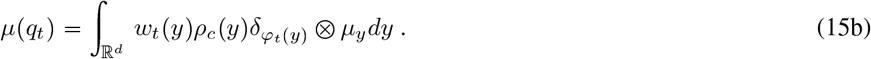

The endpoint gradient for the continuum *U* (*q*_1_), *q*_1_ = (*φ*_1_, *w*_1_) is the average of *h*_*q*_ over the feature space:

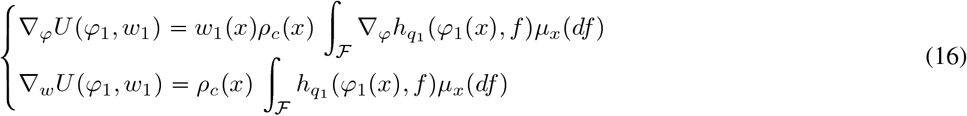

We emphasize that the varifold action gives the continuum problem unifying with LDDMM [23]; taking *I*(*y*) ∈ ℝ^+^ with 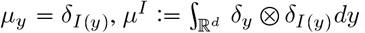, the action becomes

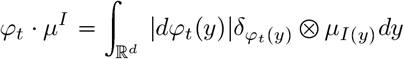

This unifies with the action of LDDMM:

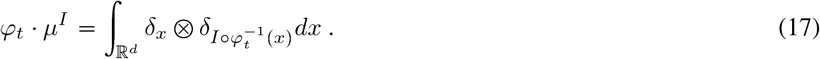

### 2.5 MRI and Digital Pathology for Tau Histology in Alzheimer’s

#### 2.5.1 Bayes Segmentation of MRI and

Figure 3 shows the multi-scale data from the clinical BIOCARD study [6] of Alzheimer’s disease within the medial temporal lobe [7, 8, 25]. The top (left) panel shows clinical magnetic resonance imaging (MRI) with the high-field 200 *μ*m MRI scale (right) shown depicting the medial temporal lobe including the collateral sulcus and lateral bank of the entorhinal cortex. Bayes classifiers for brain parcellation performs feature reduction as a key step for segmentation at tissue scales [26]. Feature reduction maps the distribution on gray levels ℱ = [0, 255] to probabilities on *N* tissue types, defined by the integration over the decision regions *θ*_*n*_ ⊂ [0, 255]:

**Figure 3:**
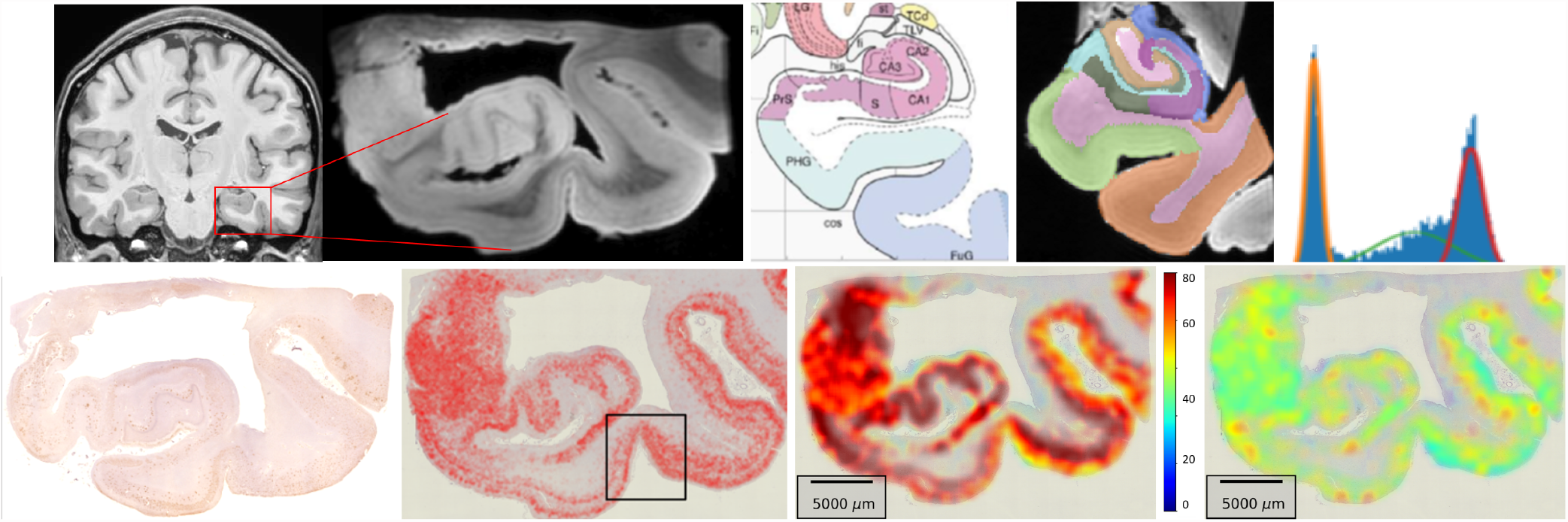
Top: Medial lobe at 1 mm and high-field 200 *μ*m MRI; Mai-Paxino atlas section of the MRI with hippocampus and entorhinal cortex; right shows the Bayes compartments. Bottom: Alzheimer 4 *μ*m Tau histology (red) from section depicted via high-field MRI (top row); right shows detected Tau particles in similar section. Box depicts trans-entorhinal region from top row. Right two panels shows mean particle size and standard deviation at *μ*m and tissue scales; deep red color denotes 80 *μ*m^2^ Tau area.

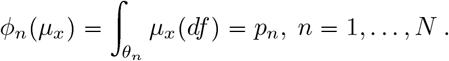

Figure 3 (top row, right) depicts a Bayes classifier for gray, white and cerebrospinal fluid compartments generated from the temporal lobe high-field MRI section corresponding to the Mai-Paxinos section (panel 3).

#### 2.5.2 Gaussian Scale-Space Resampling of Tau Histology

For histology at the molecular scales the measure encodes the detected Tau and amyloid particles 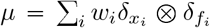 for fine scale particles with function the geometric features ℱ = ℝ^+^. Figure 3 (bottom row) shows the detected Tau particles as red dots at 4*μ*m with feature reduction done via moments on tau. We use computational lattices to interpolate between scales reapportioning particles 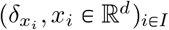 to the lattice centers 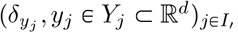, via Gaussian resampling *x* ↦ *π*(*x, Y*_*j*_) from 4*μ*m. Feature reduction maps to the first two moments at the tissue scale of mean and variance of particle size ℱ^𝓁^ ⊂ℝ^2^:

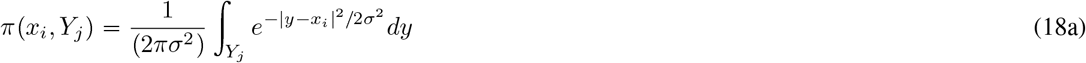

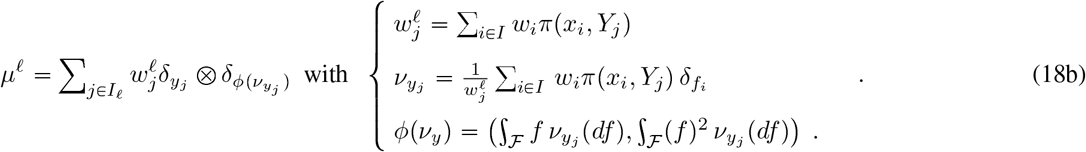

The bottom row of Figure 3 (right two panels) shows the mean and variance of the particle size reconstructed from the 4*μ*m scale: The mm scale depicts the global folding property of the tissue. The color codes the mean tissue Tau area as a function of position at the tissue scales with deep red color denoting 80 *μ*m^2^ maximum Tau area for the detected particles.

### 2.6 Cellular Neurophysiology: Neural Network Temporal Models

Single unit neurophysiology uses temporal models of spiking neurons with a “neural network” 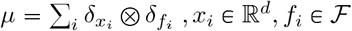 taking each neuron *x*_*i*_ modelled as a counting measure in time *N*_*i*_(*t*), *t* ⩾ 0 with the spike times the feature 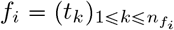:

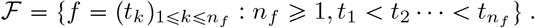

The Poisson model with intensity *λ*(*t*), *t* ⩾ *t*_0_ [10] has probabilities 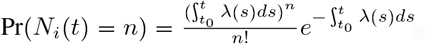.

Post-stimulus time (PST) [27] and interval histograms are used to examine the instantaneous discharge rates and inter-spike interval statistics [28]. The interval histogram abandons the requirement of maintaining the absolute phase of the signal for measuring temporal periodicity and phase locking. Synchrony in the PST is measured using binning [*b*_*i*_, *b*_*i*+1_), *i* = 1, *· · ·, B* and Fourier transforms, 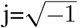:

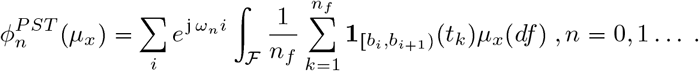

The *n* = 0 frequency computes integrated rate; each phase-locked feature is complex *ϕ*_*n*_ ∈ ℂ.

### 2.7 Scale Space Resampling of RNA to Cell and Tissue Scales

Methods in spatial-transcriptomics which have emerged for localizing and identifying cell-types via marker genes and across different cellular resolutions [4, 29–32] presents the opportunity of localizing in spatial coordinates the transcriptionally distinct cell-types. Depicted in Figure 4 are the molecular measurements at the micron scales with MERFISH [33] at three different scales.

**Figure 4:**
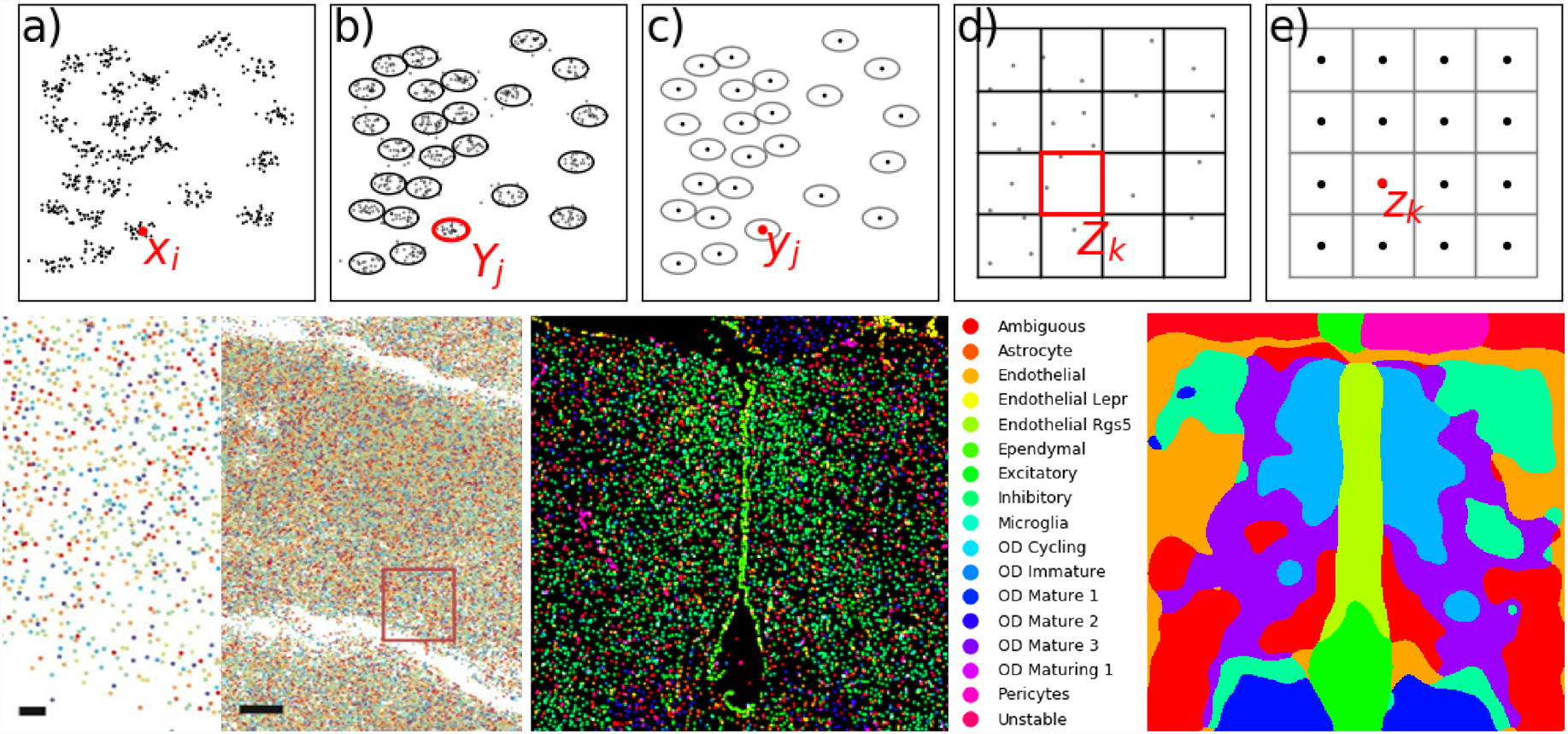
Row 1 shows cartoon of multi-scale renormalization of particles to cells and cell-centers to regular lattice representing the tissue. Row 2 shows RNA markers[33] (left) of the 167 RNA gene species (bar scales 1, 10 *μ*m); middle 17 cell types clustered on cell centers; right shows *K* = 10 means clustering to tissue with Gaussian resampling with *σ* = 25pix on the 450 × 200 pix^2^ grid.

The molecular measures represent RNA locations with sparse RNA features, 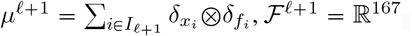. Crossing to cells 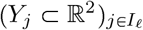 partitions into closest particle subsets defined by the distance *d*(*x*_*i*_, *Y*_*j*_) of particle *x*_*i*_ to cell *Y*_*j*_, resampling RNA particles 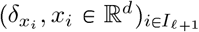 to the cell centers 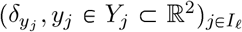 via indicator functions accumulating to nonsparse mixtures of RNA within the closest cell. The feature is the conditional probability of the 17 cell-type vector ℱ^𝓁^ ⊂ [0, 1]^17^.

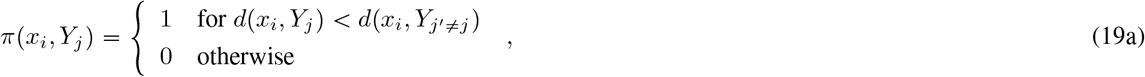

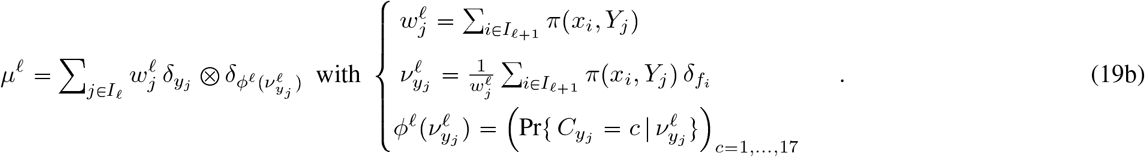

The conditional probabilities on the RNA feature vector is modelled in an independent, Gaussian expansion 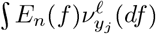 calculated via the principle components *E*_*n*_, *n* = 1, 2, ….

Resampling to tissue 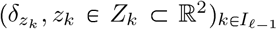 uses normal rescaling. The new feature vector becomes the probability of the cell at any position being one of 10 tissue types ℱ^𝓁-1^ ⊂ [0, 1]^10^. The probability of tissue type is calculated using 10-means clustering on the cell probabilities. The distance for 10-means clustering is computed using the Fisher-Rao metric [34] between the empirical feature laws 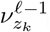. The output of 10-means are a partition of feature space ∪_*t*_ ℱ_*t*_ = ℱ^𝓁−1^ giving new features:

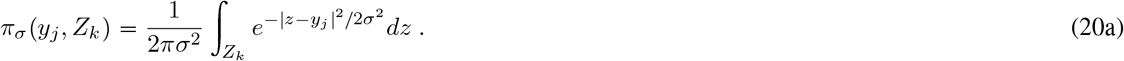

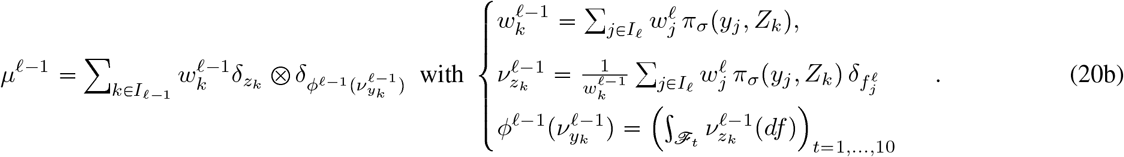

Figure 4 (bottom left panels) shows the RNA forming *μ*^𝓁+1^ depicted as colored markers corresponding to the different gene species (bar scale 1, 10 microns). Panel 2 shows the feature space of 17 cell types making up *μ*^𝓁^ associated to the maximal probability in the PCA projection from a classifier based on the mixtures of RNA at each cell location. Bottom right shows the 10 tissue features associated to the 10-means procedure. In both scales the probabilities are concentrated on one class with probability 1 and the others 0.

### 2.8 Geodesic Mapping for Spatial Transcriptomics, Histology and MRI

#### 2.8.1 Navigation Between Sections of Cell Identity in Spatial Transcriptomics

Figure 5 (top panels 1,2) shows sections (left column) from [4] depicting neuronal cell types via colors including excitatory cells eL2/3 (yellow), eL4 (orange), red eL5 (red), inhibitory cells ST (green), VIP (light blue), each classified via high dimensional gene expression features vectors via spatial transcriptomics.

**Figure 5:**
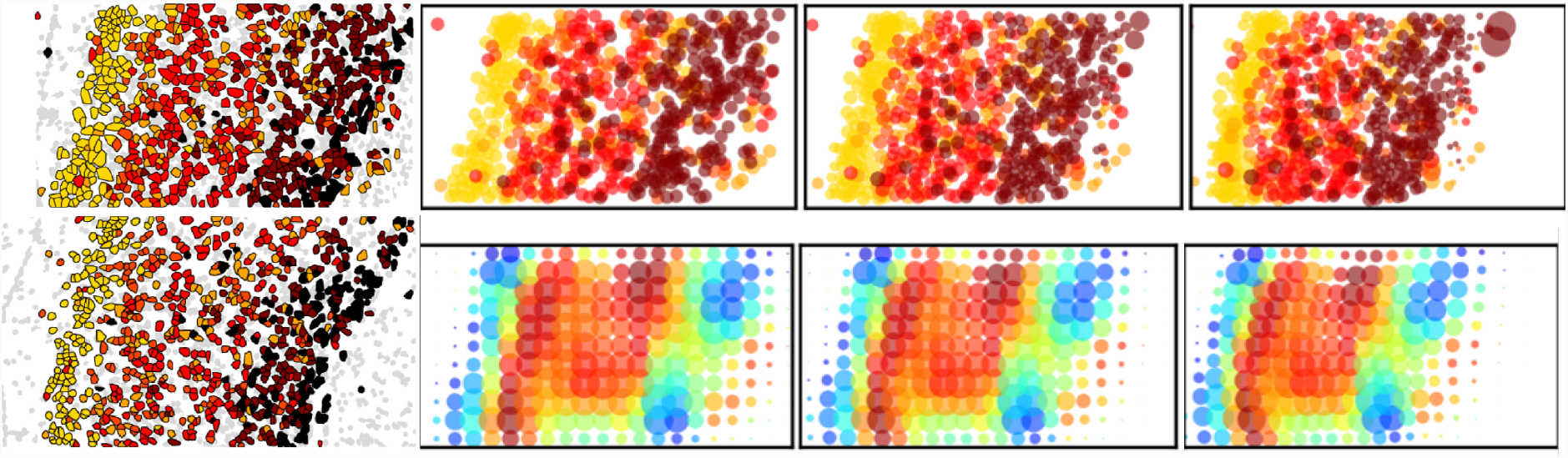
Left column: Shows neuronal cell types [4] for two sections: top template, bottom target. Right column: Panels show *φ*_*t*_ · *μ* for *t* = 0, 0.5, 1.0 of neuronal cells mapping cell types (top) and entropy (bottom). The varifold norm 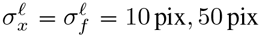 with the feature a one hot encoding of cell type. The vector fields have RKHS norm induced by the differential operator *L*_𝓁_ := ((1 − (*α*^𝓁^)^2^ ∇^2^)id_*d*_)^2^, *α*^𝓁^ = 5 pix, 50 pix with full width at half maximum (FWHM) of the Green’s kernel *FWHM* = 20 pix, 145 pix, for 𝓁 = 1, 2, respectively.

The measure 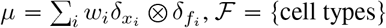 crosses to atlas scales using *π*_*σ*_ in ℝ^2^ of Eqn. (20a) with feature reduction expectations of moments, ℱ^𝓁^ = {size, variance,entropy}:

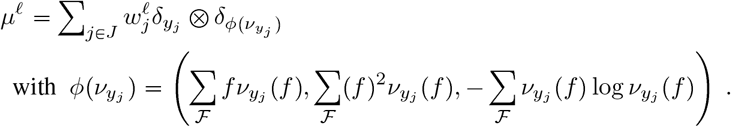

The right panels of Figure 5 shows the tissue scale features associated to the cell identity depicting and the entropy. The right panels shows the results of transforming the neuronal cells depicting the cell type (top row) and entropy feature (bottom row). The enropy is a measure of dispersion across the cell identities given by the expectation of the log probability function with zero entropy meaning the space location feature distribution *ν*_*x*_ has all its mass on 1 cell type. Geodesic mapping enforces vector field smoothness via differential operators specifying the norms in the RKHS 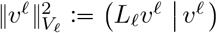 with *L*_𝓁_:= ((1 − (*α*^𝓁^)^2^∇^2^)id_*d*_)^2^, 𝓁 *<* 𝓁_*max*_.

#### 2.8.2 Navigation Between Sections of Histology

Figure 6 (rows 1 and 2) shows navigation between the cortical folds of the 4 *μ*m histology. Shown in panel 1 is a section showing the machine learning detection of the Tau particles. Columns 2,3, and 4 depict the template, mapped template and target showing the mathematical measure representation of the perirhinal cortex constructed from the positions and sizes at the 4*μ*m scale (top row) and reconstruction using Gaussian resampling onto the tissue scale (bottom row). The color codes the mean of *μ*_*x*_ representing Tau area as a function of position at the tissue scales with deep red of 80 *μ*m^2^ of Tau area the maximum value. The gradients in tau tangle area between superficial and deep layers is apparent with the deep red as high as 80 *μ*m^2^ for the innermost tissue fold. The bottom left panel shows the vector field encoding of the geodesic transformation of the of the perirhinal cortex mapping between the two sections in column 1. The narrowing of the banks of the perirhinal cortex is exhibited at the tissue scale for motions order 1000 *μ*m (brightness on scale bar).

**Figure 6:**
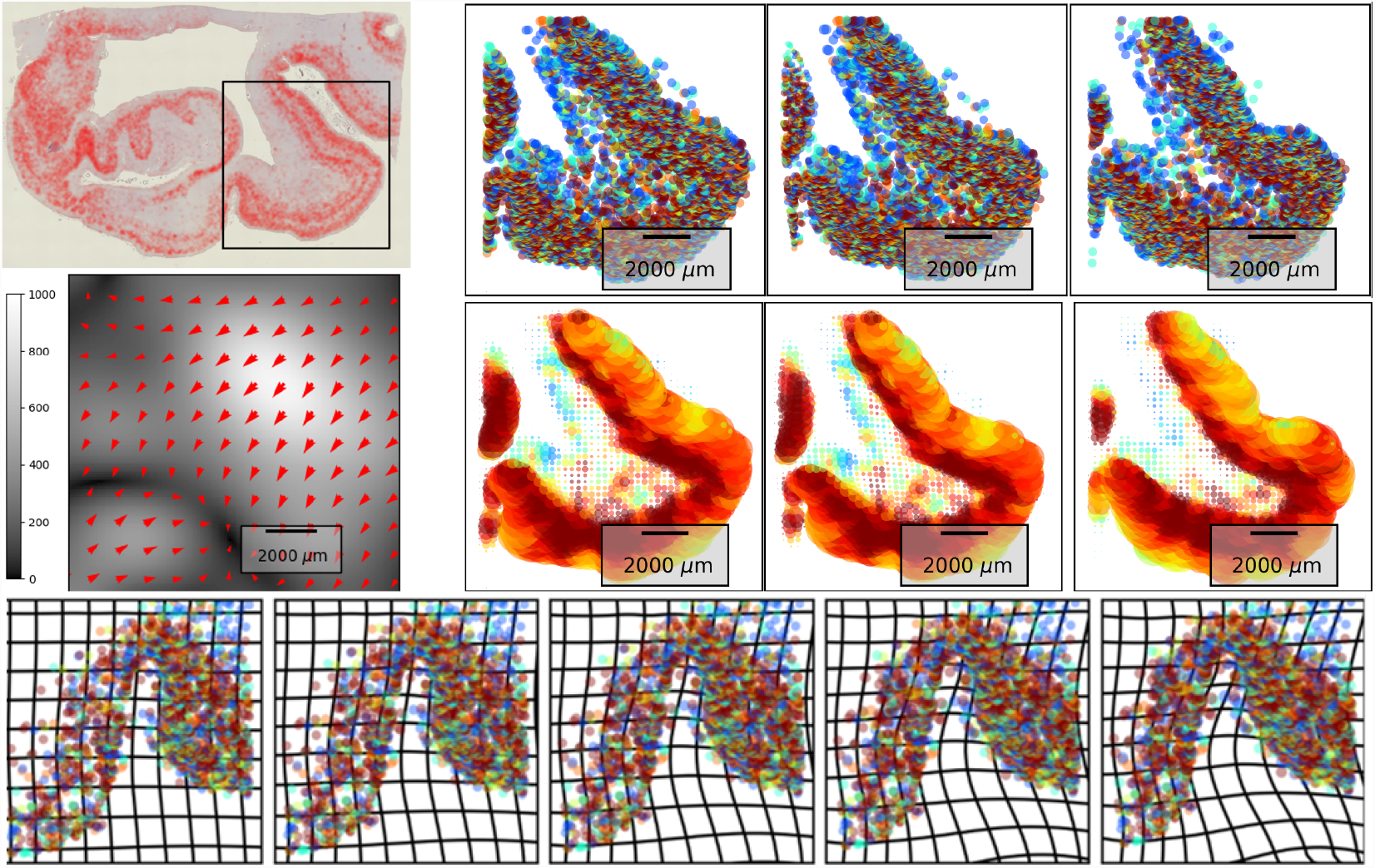
Rows 1,2: Second section 4 *μ*m histology (similar to Figure 3) with box depicting perirhinal cortex. Columns 2, 3 and 4 show template, mapped template and target at the molecular (top) and tissue scales (bottom) showing the first moments of Tau size on the perirhinal cortex; saturated red color denotes 80 *μ*m^2^ Tau area. Mapped template shows narrowing 1000 *μ*m of perirhinal sulcus. Second row left panel depicts vector field encoding of the geodesic mapping with associated scale bar. Row 3: Geodesic navigation *φ*_*t*_·*μ* of collateral sulcus (Figure 3, box) showing 1000 *μ*m widening of folds for molecular and tissue scales depicting the mean transforming. The vector field mappings have RKHS norm induced by the differential operator with *L*_𝓁_ := ((1 − (*α*^𝓁^)^2^∇^2^)id_*d*_)^2^, with *α*^𝓁^ with *FWHM* = 580*μ*m, 3300*μ*m, for 𝓁 = 1, 2 respectively. The varifold norm has 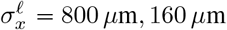 for the mappings.

Figure 6 (row 3), shows the collateral sulcus fold at the boundary of the trans-entorhinal cortex region transforming based on the normed distances between sections with deformation motions 1000 *μ*m in size. Shown is the micron scale depicting the transformation of the gyrus with the color representing the entropy of the particle identity distribution.

#### 2.8.3 Mapping Digital Pathology from Histology to MRI Scales

All of the examples thus far have created the multi-scale data generated using the resampling kernels from the finest scales. As illustrated in our early figures much of the data is inherently multi-scale, with the measurement technologies generating the coarse scale representations. Shown in Figure 7 is data illustrating our Alzheimer’s study of post mortem MR images that are simultaneously collected with amyloid and Tau pathology sections. MR images have a resolution of approximately 100 *μ*m, while pathology images have a resolution of approximately 1 *μ*m. For computational purposes the MRI template and target images were downsampled to 759 and 693 particles, respectively with the tau tangles downsampled to 1038 and 1028 particles, respectively. We treated every pixel in the MR image as a coarse scale particle with image intensity as its feature value Eqn. (17), and every detected tau tangle as a fine scale particle with a constant feature value, and performed varifold matching to align to neighboring sections. The endpoint representing the two scales is 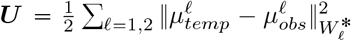. For each scale norm we use a varifold kernel given by the products of Gaussian distributions with the varifold measure norm Eqn. (22) at each scale. For the MRI scale, the weights are identically *w* = 1 with the function component given by the MRI image value; for the tau particles there is no function component making *f* = *g* with the kernel 1 for all values.

**Figure 7:**
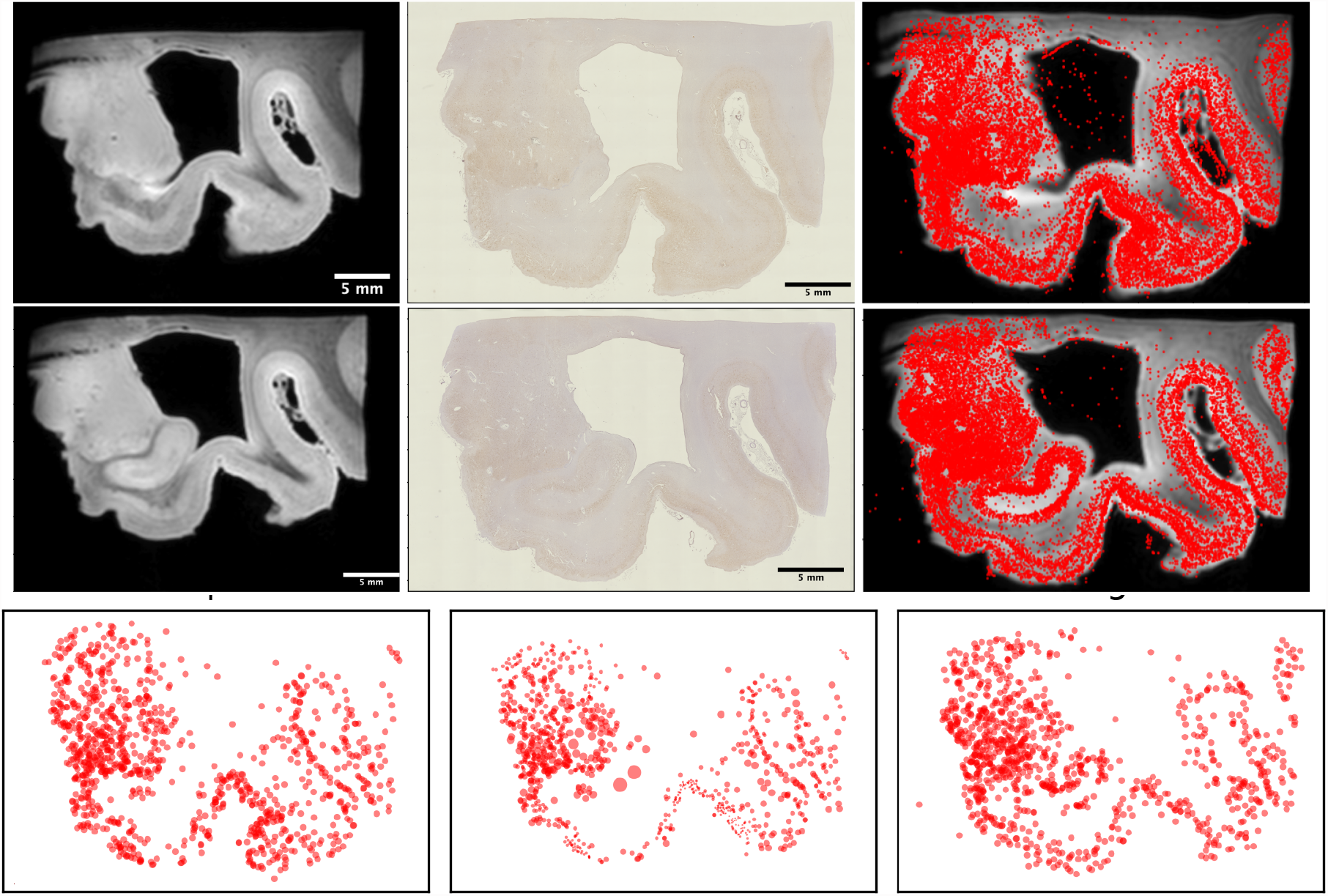
Whole brain section showing mapping MRI and histology at the multiple scales. Top two row shows the MRI and Tau histology for two sections with the detected Tau particle superimposed over the MRI (right); bottom row shows the finest scales for the template, middle the template mapped, with the right shows the target; the varifold norm has 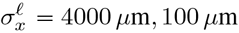. The vector field mappings have RKHS norm induced by the differential operator *L*_𝓁_:= ((1 − (*α*^𝓁^)^2^∇^2^)id_*d*_)^2^, *α*^𝓁^ giving *FWHM* = 1500 *μ*m, 6400 *μ*m, for 𝓁 = 1, 2 respectively.

The top two rows of Figure 7 shows the imaging data for both sections. The bottom row shows the transformed template image at the fine scale. The high resolution mapping carries the kernels across all the scales as indicated by the geodesic equation (14a). Notice the global motions of the high resolution of the fine particles.

## 3 Discussion

Computational anatomy was originally formulated as a mathematical orbit model for representing medical images at the tissue scales. The model generalizes linear algebra to the group action on images by the diffeomorphism group, a non-linear algebra, but one that inherits a metric structure from the group of diffeomorphisms. The formulation relies on principles of continuity of medical images as classical functions, generalizating optical flow and advection of material to diffeomorphic flow of material, the material represented by the contrast seen in the medical imaging modality such as fiber orientation for diffusion tensor imaging, and or bold contrast for gray matter content. Unifying this representation to images built at the particle and molecular biological scale has required us to move away from classical functions, to the more modern 20th century theory of non-classical generalized functions. Mathematical measures are the proper representation as they generally reflect the property that probes from molecular biology associated to disjoints sets are additive, the basic starting point of measure theory. Changing the model from a focus on groups acting on functions to groups acting on measures allows for a unified representation that has both a metric structure at the finest scales, as well as a unification with the tissue imaging scales.

The brain measure formulation, carries with it implicitly the notion of scale-space, i.e. the existence of a sequence of pairs across scales, the measure representation of the brain and the associated scale-space reproducing kernel Hilbert space of functions which correspond to the probing measurement technologies. As such part of the prescription of the theory is a method for crossing scales and carrying information from one scale to the other. Important to this approach is that at every scale we generate a new measure, therefore the recipe of introducing “measure norms” built from RKHS’s for measuring brain disparity is universal across the hierarchy allowing us to work simultaneously with common data structures and a common formalism. Interestingly, the measure norms do not require identical particle numbers across brains in brain space at the molecular scales.

The key modelling element of brain function is that the conditional feature probability is manipulated from the quantized features to the stochastic laws. These are the analogues of the Boltzman distributions generalized to the complex feature spaces representing function. As they correspond to arbitary feature spaces not necessarily Newtonian particles, we represent them simply as empirical distributions on the feature space, with the empirical measure constructed from the collapse of the fine scale to the resampled coarse scale. To model rescaling through scale-space explicitly, the two kernel transformation are used allowing us to retrieve the empirical averages represented by the determinism of the stochastic law consistent with our views of the macro tissue scales. This solves the dilemna that for the quantized atomic and micro scales cell occurence will never repeat, i.e. there is zero probability of finding a particular cell at a particular location, and conditioned on finding it once it will never be found again in the exact same location in another preparation. The properties that are stable are the probability laws with associated statistics that may transfer across organisms and species.

Importantly, our introduction of the |*dφ*(*x*)| term in the action enables the crucial property that when a tissue is extended to a larger area, the total number of its basic constituents should increase accordingly and not be conserved. This is not traditional measure transport which is mass preserving which is not a desirable feature for biological samples. Rather we have defined a new action on measures that is reminiscent of the action on *d*-dimensonal varifolds [35, 36]. We call this property “copy and paste”, the notion being that the brain is built on basic structuring elements that are conserved.

Successive refinement for the small deformation setting has been introduced in many areas associated to multigrid and basis expansions. The notion of building multi-scale representation in the large deformation LDDMM setting was originally explored F. Riesser et al. [37] in which the kernels are represented as a sum of kernels and Sommer et al. [38] in which the kernel is represented as vector bundles. In their multi-scale setting there is a post-optimization decomposition in which the contribution of the velocity field into its different components can then each be integrated. In that multi-scale setting the basic Euler-Lagrange equation termed EPDIFF remains that of LDDMM [39]. In the setting proposed here we separate the scales before optimisation via the hierarchy of layered diffeomorphisms and use a multi-scale representation of the brain hierarchy itself which is directly associated to the differomorphism at that scale. This gives the fundamental setting of the product group of diffeomorphisms with the Euler-Lagrange equation corresponding to the sequence of layered diffeomorphisms for multi-scale LDDMM [24].

The aggregation across scales from particle to tissue scales on lattices provides the essential link to inference on graphs It is natural for these aggregated features with associated conditional probability laws to become the nodes in Markov random field modelling for spatial inference; see examples in spatial transcriptomics and tissue segmentation [40]. Building neighborhood relations as conditional probabilities between lattice sites from which global probabilites laws are constructed with the Hammersley-Clifford theorem links us to Grenander’s metric pattern theory formalisms with the atoms and conditional laws 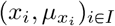 at any scale playing the roles of the generators.

## 4 Materials and Methods

### 4.1 Experimental and Technical Design

The objective of this research is to unify the molecular representations of spatial transcriptomics and cellular scale histology with the tissue scales of Computational Anatomy for brain mapping. To accomplish this we designed a mathematical framework for representing data at multiple scales using generalized functions, and mapping data using geodesic flows of multiple diffeomorphisms. We illustrate the method using several examples from human MRI and digital pathology, as well as mouse spatial transcriptomics.

### 4.2 Computational Lattices for Interpolating Brain Measures

We use computational lattices 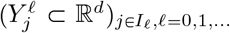 to interpolate 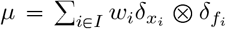 onto *μ*^𝓁^ at scale 𝓁. Then 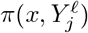 is chosen to share its probability concentrated to the lattice cell centers 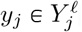:

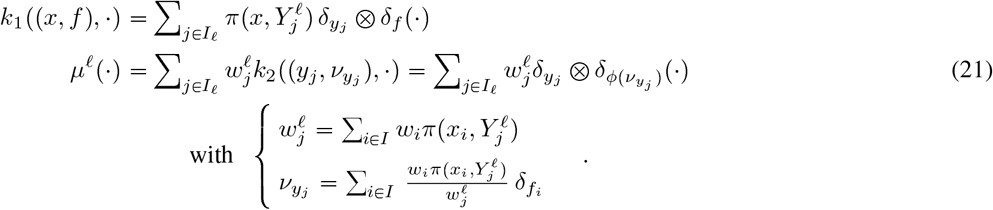

### 4.3 Gaussian Kernel Varifold Norm

Our varifold norm construction models the measures as elements of a Hilbert space *W* * which is dual to an RKHS *W* with a kernel *K*_*W*_. We introduce the dual bracket notation for *h* ∈ *W, μ* ∈ *W* *, 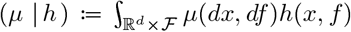. The norms are generated by integrating against the kernel according to (10) written with the dual bracket 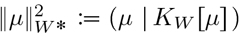; the multi-scale norm is given by 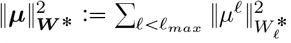.

To ensure the brain measures are elements of *W* * dual to the RKHS *W*, the kernel *K*_*W*_ is chosen to densely and continuously embed in bounded continuous functions 𝒞_*b*_(ℝ^*d*^ × ℱ, ℝ) so that the signed measure spaces ℳ_*s*_(ℝ^*d*^ *×*ℱ) of brains are continuously embedded in the dual spaces 𝒞_*b*_(ℝ^*d*^ × ℱ, ℝ)*. An example is the Gaussian kernel (22) which satisfies this condition, the kernel taken as separable Gaussians with | · | Euclidean distance:

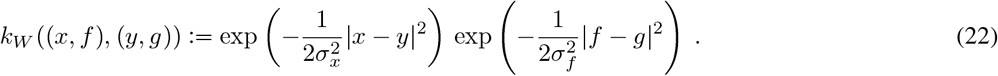

For measures 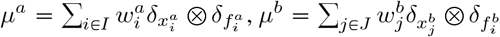 gives

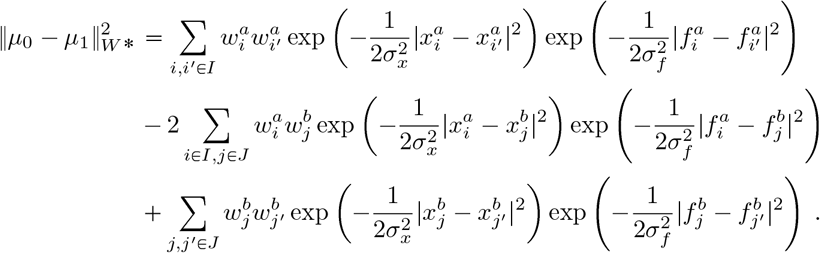

For data carrying position information but no feature values (such as tau tangle locations), each *f*_*i*_, *f*_*j*_ is constant and the resulting exponential terms are all 1.

For data carrying position information but no feature values (such as tau tangle locations), each *f*_*i*_, *f*_*j*_ is constant and the resulting exponential terms are all 1.

### 4.4 The Riemannian Distance Metric on the Hierarchical Group

The diffeomorphism group acts on the hierarchy ***φ***·***µ*** component-wise Eqn. (7b) with the multi-scale group the product

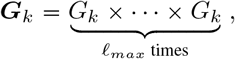

with elements ***φ*** ∈ ***G***_*k*_ satisfying the law of composition component-wise 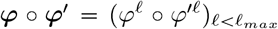. The group *G*_*k*_ supporting *k*-derivatives of the diffeomorphisms builds from 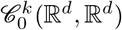 space of k-times continuously differentiable vector fields vanishing at infinity and its partial derivatives of order *p* ⩽ *k* intersecting with diffeomorphisms with 1-derivative:

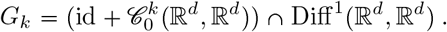

Dynamics occurs via group action generated as a dynamical system in which the multi-scale control 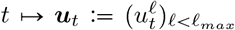 flows the hierarchy *t* ↦ ***φ***_*t*_ satisfying 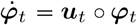 of (8a). The control is in the product 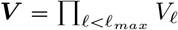, each space an RKHS with norm-square 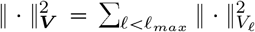 selected to control the smoothness of the vector fields. The hierarchy of spaces are organized as a sequence of continuous embeddings:

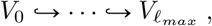

where 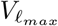 is an additional layer containing the others with 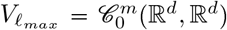 defined as a space of m-times continuously differentiable vector fields vanishing at infinity as well all its partial derivatives of order *p* ⩽ *m*.

The hierarchy is connected via successive refinements *u*^𝓁^ = *u*^𝓁 −1^ + *v*^𝓁^, *u*^0^ = *v*^0^ expressed via the continuous linear operator ***A*** : ***V*** → ***V*** with ***v*** = ***Au***. The control process (***u***_*t*_)_0⩽*t*⩽1_ ∈ *L*^2^([0, 1], ***V***) has finite square-integral with total energy

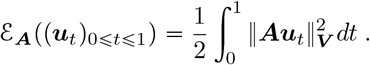

Optimal curves which minimize the integrated energy ℰ_***A***_ between any two fixed boundary conditions (BC) ***φ***_0_ = **Id** and ***φ***_1_ which is accessible with a path of finite energy extends the LDDMM setting [23] to a hierarchy of diffeomorphisms and describes a geodesic for an associated Riemannian metric and multi-scale LDDMM [24] on ***G***_*k*_:

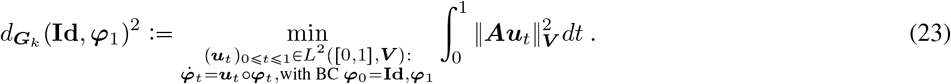

Existence of solutions for minimizers over ***u*** of (23) when 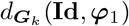 is finite can be established when *m*⩾ *k*⩾ 1.

### 4.5 Geodesic Multi-Scale LDDMM via Hamiltonian Control

The Hamiltonian method reduces the parameterization of the vector field to the dynamics of the particles that encode the flow of states (12). We write the dynamics explicitly as a linear function of the control, which defines the flow of the measures:

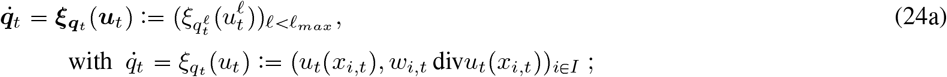

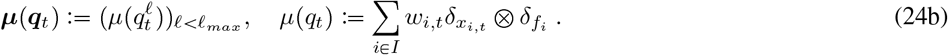

The control problem satisfying (11) reparameterized in the states becomes, for *α* > 0:

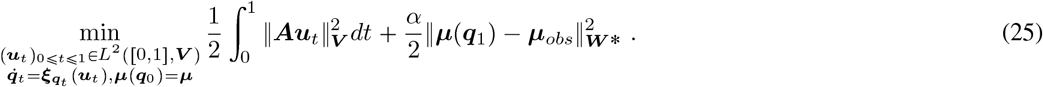

Hamiltonian control introduces the co-states 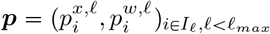 via the Hamiltonian

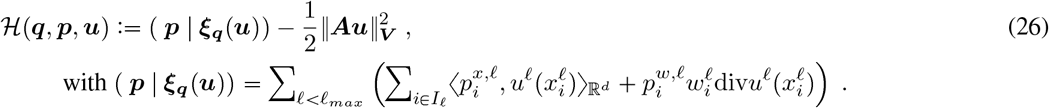

Under the assumption 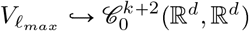 the Pontryagin maximum [21] gives the optimal control satisfying for every 𝓁:

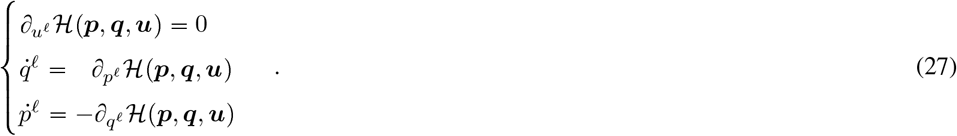

The shape of Brainspace is given by its geodesics.

**Statement 1** (Geodesics of Space). *Assume that* 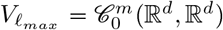 *with m* ⩾*k* + 2. *If* (***u***_*t*_)_0⩽*t*⩽1_ *is a solution of the optimal control problem* (25) *then there exists time-dependent co-state* (*t* ↦ ***p***_*t*_) *for any i ∈I*_𝓁_ *for* 𝓁*all satisfying (proof Appendix A):*

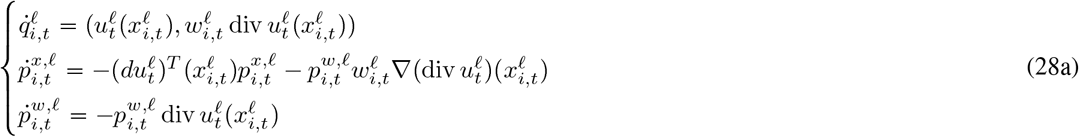

*The optimal control satisfies* 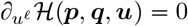 *and* ***v*** = ***Au***, *for any* 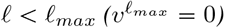:

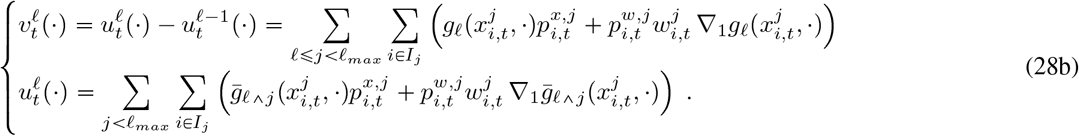

**Statement 2** (Integral equations for Hamiltonian-Momentum). *Assuming* 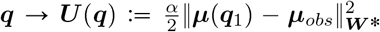 *is* 𝒞 ^1^ *in* ***q***, *the geodesic co-state flowing from t* = 0, 1 *satisfy (proof Appendix B):*

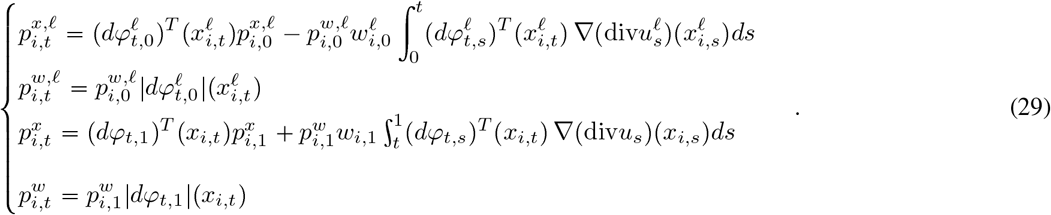

### 4.6 Gradients of the Endpoint Varifold Matching Norm

The gradients of (14c) are efficiently rewritten using the state *q*_*t*_ = (*x*_*i,t*_, *w*_*i,t*_)_*i* ∈*I*_ to define the norm-square in terms of *h*_*q*_ continuously differentiable in *x* and bounded 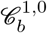 giving the smooth endpoint term determining the smooth gradients:

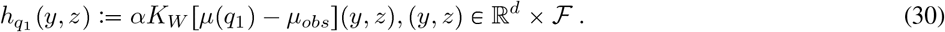

We take the variation 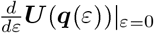 varying each term 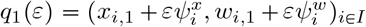 with dependence on 𝓁-scale implied.

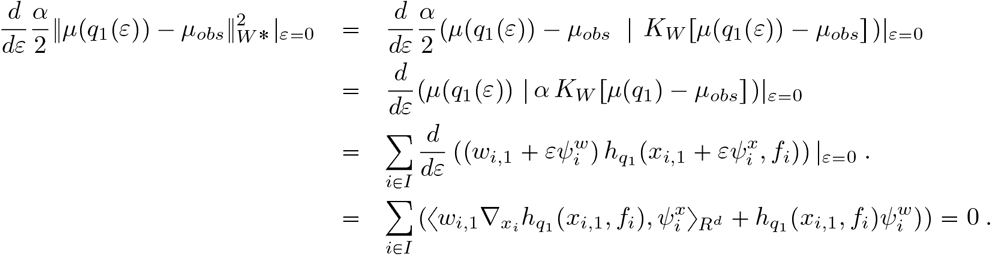

These represent the gradients of (14c).

The tissue continuum has 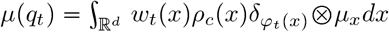 with 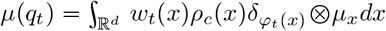 and *q*_*t*_ := (*φ*_*t*_, *w*_*t*_ = *w*_0_|*dφ*_*t*_1). The average of 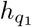 over the feature space determines the boundary term variation.

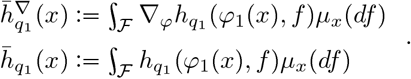

With *q*_1_ = (*φ*_1_, *w*_1_) take the variation *φ*_1_ →, *φ*_1_(*ε*) = *φ*_1_ + *εψ*^*φ*^, *w*_1_ →, *w*_1_(*ε*) = *w*_1_ + *εψ*^*w*^:

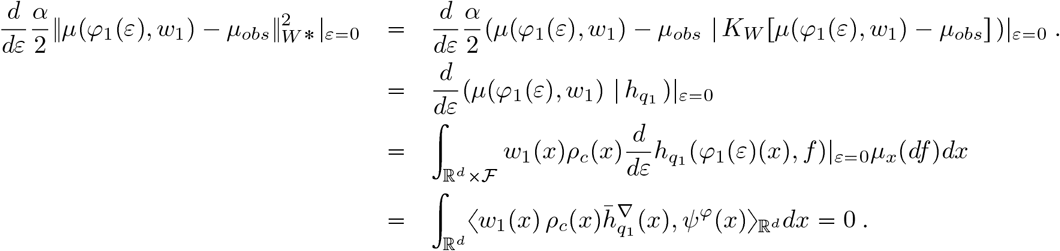

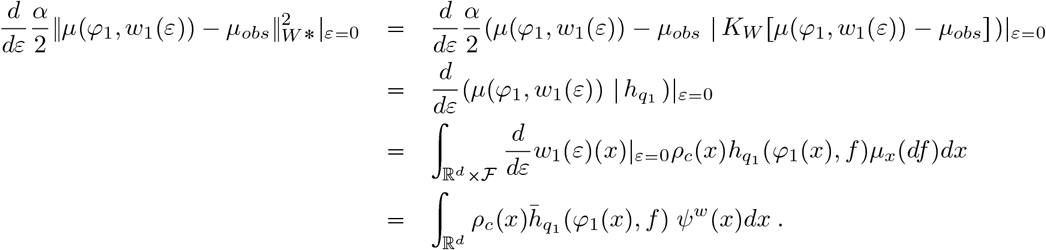

This gives the gradients of the matching endpoint Eqn. (16). We note that computing the variation 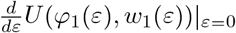 requires *U* as a function of *φ* is 𝒞 ^1^ for 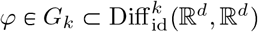 when *k* ⩾ 2.

## Author Contributions

MM and AT designed the theoretical framework and DT built the computational implementation. All authors contributed to writing and editing the manuscript.

## Data Availability

The major contribution of this work is a mathematical and computational framework for modeling hierarchical neuroimaging data. Specific datasets shown in examples are for illustrative purposes, but can be made available upon request.

## A Hamiltonian Control Statement

We take for our RKHS *V*_𝓁_, 𝓁*<*𝓁_*max*_ with the operators *L*_𝓁_ : *V*_𝓁_ → *V*_𝓁_* defining the isometries such that the inner products 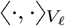 satisfy for any 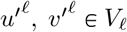:

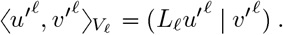

## Appendix Statement 1.

*We assume that* 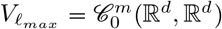 *with m* ⩾ *k* + 2. *If* ***u***. *is a solution of the optimal control problem* (25) *then there exists time-dependent co-state* (*t* ↦ ***p***_*t*_) *such that for any i* ∈ *I*_𝓁_ *for all* 𝓁 :

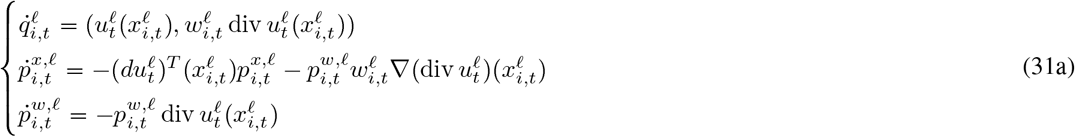

*The optimal control satisfies* 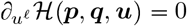 *and* ***v*** = ***Au***, *for any* 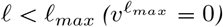:

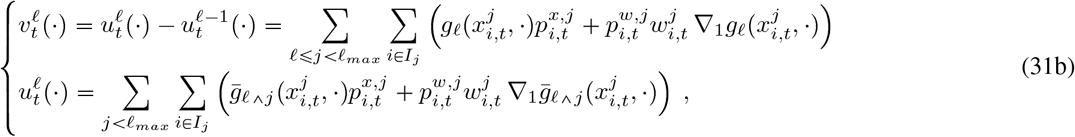

*with* 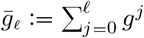 *and* ∇_1_*g*(*a, b*) *denotes the gradient of g*_𝓁_ *with respect to the first variable a*.

*Proof*. Under the assumption 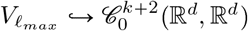 then we have (***u, q***) ↦ ***ξ***_***q***_(***u***) is 𝒞 ^2^ and standard results of optimal control theory applying the Poyntryagin maximum principle [21] gives

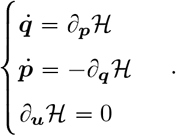

Taking the variation for 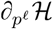 of the Hamiltonian for scale 𝓁 varies 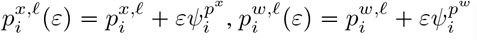 :

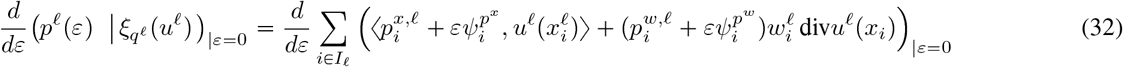

which gives the first equation for the state velocity (31a).

Taking the variation for 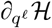 of the Hamiltonian for scale 𝓁 varies 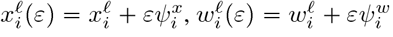:

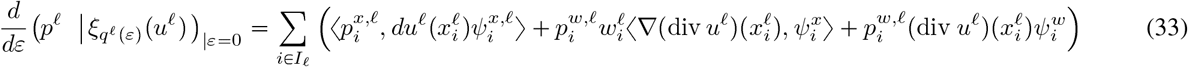

so that we get the second two equations of (31a). To calculate 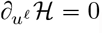, define *u*^𝓁^(*ε*) = *u*^𝓁^ + *εψ*^*u*^ implying *v*^𝓁+1^(*ε*) = *v*^𝓁+1^ − *εψ*^*u*^, *v*^𝓁^(*ε*) = *v*^𝓁^ + *εψ*^*u*^ for *ψ*^*u*^ ∈ *V*_𝓁_. We have

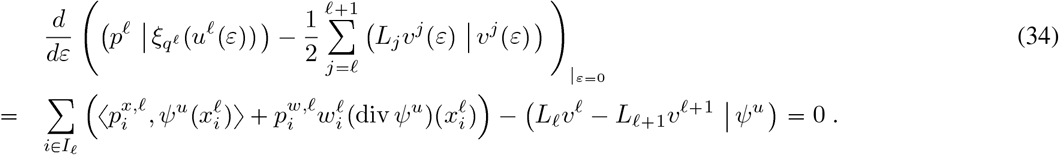

After summation of (34) for 𝓁 ⩾ 𝓁_′_, we get for any *ψ*^*u*^ ∈ *V*_𝓁′_ that

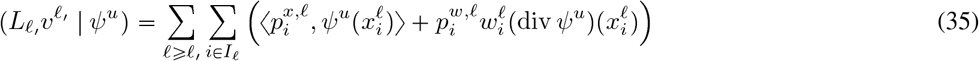

Now, for any *x, α* ∈ ℝ^*d*^, consider 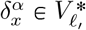 such that 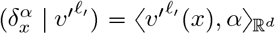 for any 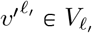. The reproducing property on 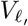 gives 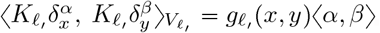. We get from (35) for 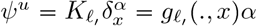 that

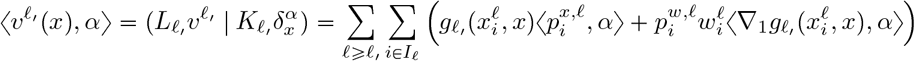

so that we get the first equality above of (31b) for 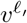 for any 𝓁_′_ < 𝓁_*max*_.

Now since 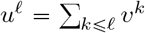 we deduce the equality for *u*^𝓁^, 𝓁 < 𝓁_*max*_ given in (31b).□

### B Hamiltonian co-state momentum integrable dynamics

We omit the superscripts 𝓁 below since co-states and states and flows are at the same scale.

## Appendix Statement 2.

*Assume* ***q*** → ***U*** (***q***) *is* 𝒞^1^ *in* ***q***, *then the co-state integral equations* (14b) *flowing from t* = 1 *solves the Hamiltonian differential equations* 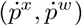 *of* (31a); *the integral equations flowing from t* = 0 *satisfy:*

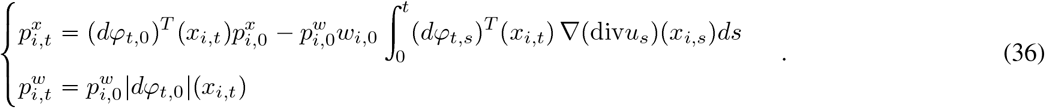

*Proof*. First take 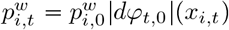 and show it satisfies (31a). For *w*_*i,t*_ = *w*_*i*_|*dφ*_*t*_| (*x*_*i*_), *x*_*i,t*_ = *φ*_*t*_(*x*_*i*_) then

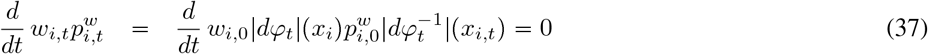

which implies 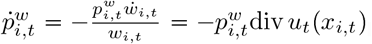.

The 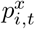 satisfies (31a); rewrite the integral solution using 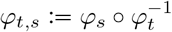 and the identities.

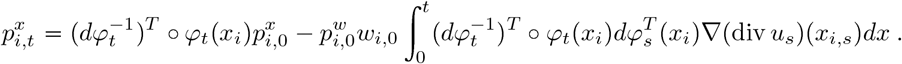

Differentiating requires the identity 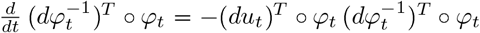 (see below):

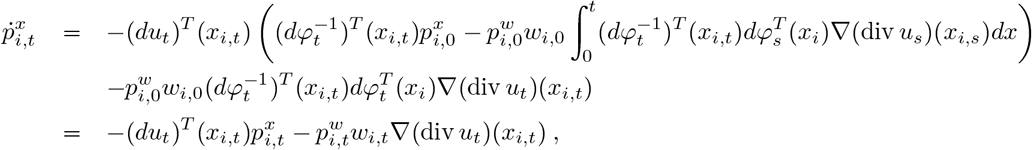

where the last equality uses (37), 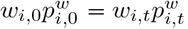. The remaining identity follows 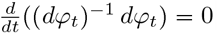:

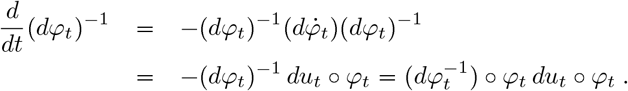

The optimal control (14a) is written in the endpoint 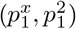 with the boundary conditions (14b). Using the form of 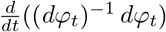 gives 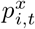 in the boundary *t* = 1 satisfies the differential equation. For 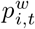 then constancy 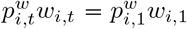 with 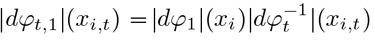 gives the endpoint:

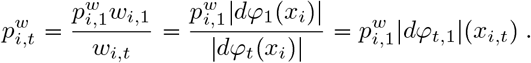

□

